# Recently emerged and diverse lineages of *Xanthomonas perforans* have independently evolved through plasmid acquisition and homologous recombination originating from multiple *Xanthomonas* species

**DOI:** 10.1101/681619

**Authors:** E. A. Newberry, R. Bhandari, G.V. Minsavage, S. Timilsina, M. Jibrin, J. Kemble, E. Sikora, J.B. Jones, N. Potnis

**Affiliations:** Department of Entomology and Plant Pathology, Auburn University, AL 36849; Department of Plant Pathology, University of Florida, FL 32611

## Abstract

*Xanthomonas perforans* is the predominant pathogen responsible for bacterial leaf spot of tomato and *X. euvesicatoria* of pepper in the southeast United States. Previous studies have indicated significant changes in the *X. perforans* population collected from Florida tomato fields over the span of two decades including a shift in race, diversification into three genetic groups, and host range expansion to pepper. Recombination originating from *X. euvesicatoria* was identified as the primary factor driving the diversification of *X. perforans* in Florida. The aim of this study was to genetically characterize *X. perforans* strains that were isolated from tomato and pepper plants grown in Alabama and compare them to the previously published genomes available from GenBank. Surprisingly, a maximum likelihood phylogeny coupled with a Bayesian analysis of population structure revealed the presence of two novel genetic groups in Alabama, which each harbored a different transcription activation-like effector (TALE). While one TALE, *avrHah1*, was associated with adaptation of *X. perforans* to pepper, the other was identified as a new class within the *avrBs3* family, designated here as *pthXp1*. Examination of patterns of homologous recombination between *X. perforans* and other closely related *Xanthomonas* spp. indicated that the lineages identified here emerged in part through recent recombination events originating from xanthomonads associated with diverse hosts of isolation. Our results also suggest that the evolution of pathogenicity to pepper has likely emerged independently within *X. perforans* and in one lineage, was associated with the recombination-mediated remodeling of the Xps type II secretion and TonB transduction systems.

**Importance:** The emergence of novel pathogen lineages has important implications in the sustainability of genetic resistance as a disease management tool in agricultural ecosystems. In this study, we identified two novel lineages of *X. perforans* in Alabama. While one lineage was isolated from symptomatic pepper plants, confirming the host range expansion of *X. perforans*, the other lineage was isolated from tomato and acquired a novel transcription activation-like effector, *pthXp1*. Unlike AvrBs4, PthXp1overcomes Bs4-mediated resistance in tomato, indicating the evolution of this novel lineage towards fitness on this host. Our findings also show that different phylogenetic groups of the pathogen have experienced independent recombination events originating from multiple *Xanthomonas* species. This suggests a continuous gene flux between related xanthomonads associated with diverse plant hosts which results in the emergence of novel pathogen lineages and associated phenotypes, including host range expansion.

## Introduction

The importance of genetic exchange in bacterial evolution can be traced back as early as to the 1960’s, where plasmid-mediated transfer of penicillin resistance was documented among members of the *Enterobacteriaceae* (1). With the increasing availability of bacterial genomes, it has since become clear that the exchange of mobile genetic elements including plasmids, bacteriophage, genomic islands, and other mechanisms of horizontal gene transfer are commonplace among bacterial populations. Such genetic exchanges can confer traits imparting phenotypic and genotypic plasticity to a bacterial species in response to changes in the environment (2), thus, facilitating adaptive evolution. In addition to horizontal gene transfer, many bacteria undergo homologous recombination. This is a process similar to meiotic recombination, in which segments of a bacterial genome are replaced by homologous sequences from a donor organism, resulting in a mosaic pattern of loci with distinct evolutionary histories (3).

In the absence of barriers such as adaptive incompatibility, related bacterial lineages are expected to display evidence of admixture in their evolutionary history when they inhabit overlapping environmental niches (4). Therefore, patterns of recombination are hypothesized to reflect a corresponding microbial ecology and maintain the cohesion of various bacterial species as monophyletic groups (5). In contrast to many genera of plant-pathogenic bacteria including *Pseudomonas*, *Ralstonia*, and *Burkholderia*, among others which are abundant in multiple environments including water and soil, the life history of *Xanthomonas* spp. has traditionally been considered to be restricted to plants (6). Most xanthomonads exhibit a high degree of host-specificity, with the individual species (or the pathovars found within them) each forming a genetically monomorphic cluster of lineages. This evolutionary trend suggests that the divergence of *Xanthomonas* species/pathovars is primarily driven by ecological isolation associated with adaptation to a particular host (7–9).

Among the numerous plant diseases caused by xanthomonads, bacterial leaf spot of tomato (*Solanum lycopersicum*) and pepper (*Capsicum annuum*) is atypical as four distinct species including *X. gardneri, X. euvesicatoria, X. perforans*, and *X. vesicatoria* have converged on these hosts to cause the same disease (10). These species may differ in geographic distribution as well as in the molecular mechanisms employed in pathogenesis (11–13). Likewise, temporal shifts in the species and pathogen races (as determined by gene for gene interactions) responsible for the disease in specific tomato and pepper production regions have been documented over the years (12, 14). Although these fluxes in the pathogen population are subject to random genetic drift associated with patterns of international trade, evolutionary and ecological factors such as horizontal gene transfer, homologous recombination, and interspecific competition via the production of antagonistic bacteriocins are also significant factors contributing to these population dynamics (15–19). Therefore, bacterial leaf spot presents an attractive model system enabling the investigation of adaptive evolution in a bacterial population mediated by recombination, host selection pressure, and interspecific competition.

Currently, *X. perforans* can be divided into three different phylogenetic groups (19). Until the isolation of a single *X. perforans* strain from an infected pepper sample during the 2010 season in Florida (20), the host-range of this bacterial pathogen was thought to be restricted to tomato. This observation led to subsequent investigations which revealed that in the absence of effector triggered immunity induced by a single avirulence gene (either *avrBsT* or *avrXv3*), *X. perforans* strains representative of the three phylogenetic groups differed in their ability to multiply and cause disease when infiltrated into pepper leaves (20). Intriguingly, genomes of the pepper-pathogenic, group 2 *X. perforans* strains displayed evidence of extensive recent recombination originating from *X. euvesicatoria* (which is the predominant pathogen of pepper), whereas similar signatures were reported to be minimal among the group 1, tomato-limited strains (13, 19).

Given the close evolutionary relationship (average nucleotide identity values greater > 98%) and overlapping host-range of *X. perforans* and *X. euvesicatoria*, it is not surprising to find evidence of genetic exchange between them. In fact, several authors have proposed that they be considered pathovars of *X. euvesicatoria*, rather than distinct species (21, 22) and are closely related to a number of strains located within the *X. euvesicatoria* species complex, *sensu* Parkinson et al. (23). The strains that belong to this larger phylogenetic group were classified into several species including *X. alfalfae*, *X. axonopodis*, *X. perforans* and *X. euvesicatoria* and are associated with diseases of diverse monocot and dicotyledonous plant families (23). While the importance of homologous recombination in facilitating the emergence of novel *X. perforans* lineages has been established (18, 19), the extent of genetic exchange across the larger *X. euvesicatoria* species complex and specific functional pathways affected by homologous recombination remains to be explored.

A detailed knowledge of the population structure of xanthomonads responsible for bacterial leaf spot of tomato and pepper in the southeast United States comes primarily from surveys conducted in Florida and to a limited extent in Georgia, South Carolina and North Carolina, where to date, only a single *X. perforans* strain has been isolated from naturally infected pepper plants (20). In a recent survey of the *Xanthomonas* population responsible for the disease in Alabama, we were readily able to isolate *X. perforans* strains from pepper plants grown in several Alabama counties (Potnis et al., *unpublished data*). This indicates a recent shift in the host range of the pathogen and an emerging threat to pepper production. Here, we sequenced the genomes of eight *X. perforans* strains that were isolated from tomatoes and peppers grown in Alabama and compared them with previously published genome data available from GenBank. Using a Bayesian statistical approach, we refined the population structure of *X. perforans* with the identification of two novel phylogenetic groups. Our findings indicate that the recent emergence of these genetic groups was associated with the acquisition of novel transcription activator-like effectors and independent recombination events originating from multiple species found within the *X. euvesicatoria* species complex.

## Results

### Reconstruction of the *X. perforans* (*Xp*) core genome reveals the presence of two novel genetic clusters composed of *Xp* strains collected in Alabama

A maximum likelihood phylogeny constructed from a concatenated alignment of 16,501 high quality, core-genome single nucleotide polymorphisms (SNPs), coupled with a Bayesian analysis of population structure (24), revealed the presence of six distinct genetic clusters within *X. perforans* (Fig. 1). Sequence clusters (SCs) 1 through 4 corresponded to the previously described population structure of *X. perforans* strains collected in Florida (19, 20), while SCs 5 and 6 were composed exclusively of Alabama strains sequenced in this study. Each of the genetic clusters inferred in this analysis were primarily clonal within the lineage, except for SC2 (referred to as Group 1b by Schwartz et al. (20), which exhibited diversity. SC3 (Group 2) contained the Florida strain originally isolated from pepper (Xp2010) as well as other strains demonstrated as pathogenic to this host under artificial inoculation conditions (20). A single Alabama strain (ALS7E) isolated from pepper clustered within this group, while the remaining pepper strains (ALS7B, AL65, and AL66) and one tomato strain (AL1) formed a novel phylogenetic lineage, designated here as SC6. The Alabama strain AL57 grouped with other *X. perforans* strains collected in Florida within SC4 (Group 3). This genetic cluster branched from SC5, which was composed of two Alabama strains (AL33 and AL37) isolated from tomato plants.

**Figure 1.**
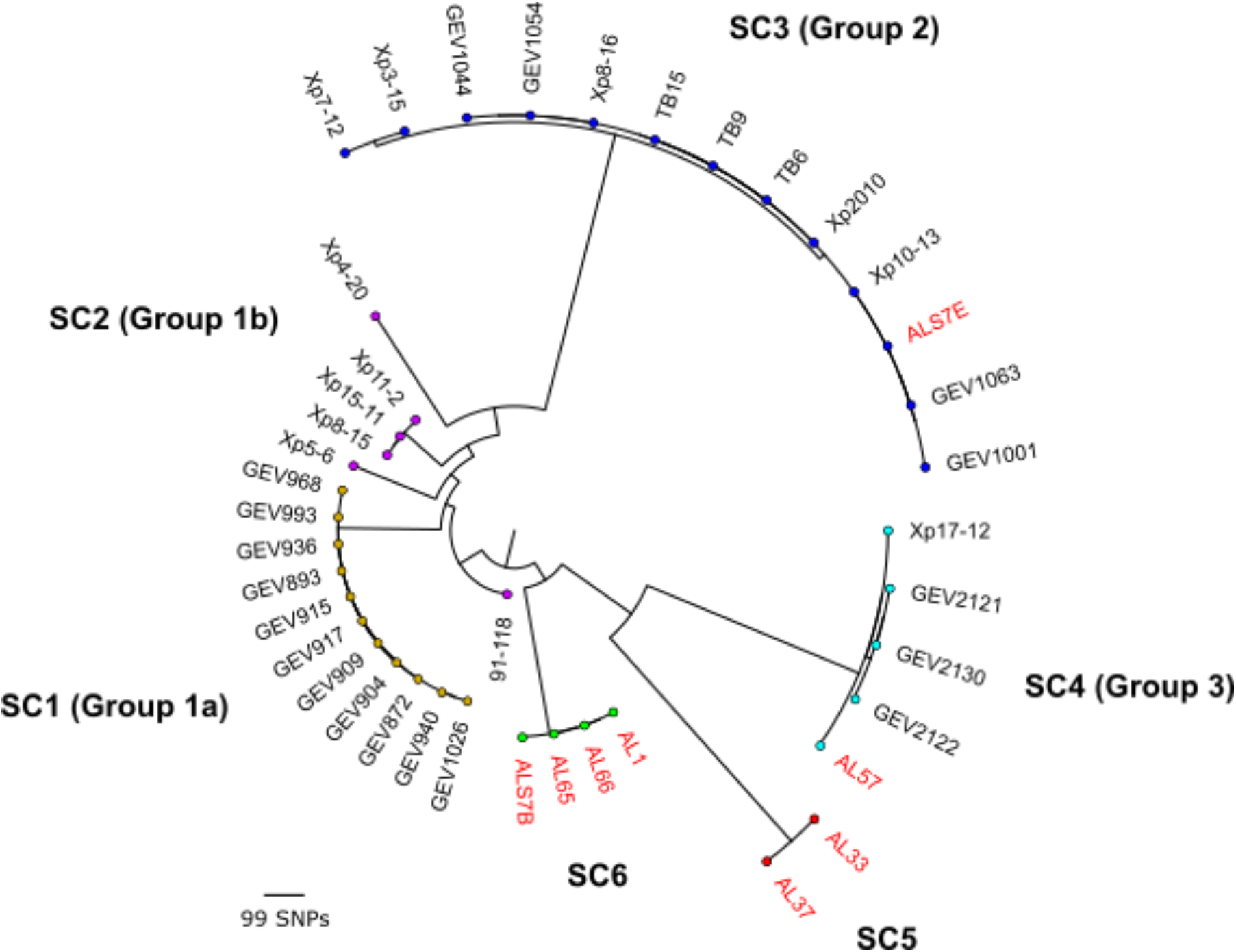
Mid-point rooted, maximum-likelihood phylogeny of 41 *X. perforans* strains isolated tomato and pepper plants grown in Florida and Alabama based on a concatenated alignment of 16,806 core-genome single nucleotide polymorphisms (SNPs). The tips are color-coded according to the sequence clusters (SCs) identified in first level of HierBAPS hierarchy (24). The phylogroup designations of strains described previously (19, 20) are shown in parentheses and strains sequenced in this study are highlighted in red. The scale bar indicates the number of substitutions per site.

### Analysis of type 3 secreted effector (T3SE) repertoires provides evidence for the plasmid mediated acquisition of several accessory T3SEs, including the transcription activation-like effector (TALE) *avrHah1*, among *Xp* strains collected in Alabama

A total of 24 T3SEs were conserved among the *Xp* strains sequenced in this study (Table 2). These results were largely concordant with conserved effectors identified across the *Xp* population characterized in Florida (20), except for *xopE2*, which was carried by only three strains (AL33, AL37, and ALS7E) sequenced here and was present in contigs which displayed 100% identity to plasmid pLH3.2 (NZ_CP018476.1) from *Xp* strain LH3. These contigs also contained genes encoding for copper resistance including *copA*, *copB*, *copF*. Strains AL33, AL37, and AL57 carried an intact copy of the avirulence gene *avrXv3*, which was disrupted by an insertion sequence in the other Alabama strains. The inability of the latter strains to induce an HR in the pepper cv. Early Cal Wonder (ECW) confirmed the non-functionality of *avrXv3*, while the former stains with the intact allele elicited an HR in ECW (Table 2).

Several other variable T3SEs were identified among the Alabama strains along with evidence of their presence in putative plasmids. The T3SEs *xopAQ* and disrupted copy of *xopE3* were present in strains AL37 and AL57, while absent from the genome assemblies of other Alabama strains. These effectors were present in the same contig assembled by plasmidSPAdes in strain AL57 which displayed significant homology (99% nucleotide identity and 94% query coverage) to an unnamed plasmid found in *X. campestris* pv. *campestris* strain CN18 (CP017322.1). Likewise, an unidentified T3SE with homology to a*vrBsT* (71% aa identity) was also carried on a putative plasmid in the same two strains. The top BLAST hit for these contigs (94% nucleotide identity over 87% query coverage) was plasmid pXCARECAE29 (NZ_CP034654.1) from *X. campestris* pv. *arecae* strain NCPPB 2649.

Analysis with plasmidSPAdes software also revealed the presence of several contigs in all but one of the strains (ALS7B) located in genetic cluster SC6, which displayed 100% nucleotide identity and over 99% query coverage to the completed plasmid pJS749-3.2 (NZ_CP018730.1) from *X. gardneri* strain JS749-3. The sequences corresponding to this putative plasmid were divided between two and five contigs per strain and summed from 43 to 45 kb, which was consistent with the size pJS749-3.2 (46 kb). Analysis of these contigs with tBLASTn produced significant hits for the TALE *avrHah1* and the T3SE *xopAO*. The activity of a*vrHah1* among these strains was supported through inoculations in the pepper cv. ECW, which produced the profuse water-soaking phenotype associated with this TALE (Fig. S1). Further evidence was obtained through inoculations in pepper cv. ECW30R, which resulted in a hypersensitive resistance (Table 2). The contigs assembled by plasmidSPAdes and BLAST search results are presented in Table S2.

### Identification of a novel class of TALE within the *avrBs3* family among different genetic backgrounds of *Xp*

Screening of the genome assemblies for T3SEs with tBLASTn indicated the presence of a putative TALE with homology to *avrBs3* in strains AL57 and AL37 of SCs 4 and 5 respectively. Phusion polymerase chain reaction (PCR) utilizing primers designed to anneal to conserved loci within the N- and C-terminal domains of the *avrBs3* effector family produced an ~2.9 kb amplicon (Fig S2). Sanger sequencing of the amplicon utilizing internal primers flanking the repeat region of *avrBs3* revealed that the TALE was comprised of 15.5 tandem repeats, each 102 bp in length. A BLAST search of the NCBI non-redundant protein database using the N- and C-terminal domains of the protein displayed 99 and 100% identity, respectively, to the *avrBs3* allele (X16130.1) found in *X. vesicatoria*.

Despite this close homology to *avrBs3*, an alignment of the repeat variable di-residue (RVD) sequences with AnnoTALE software indicated that this gene could not be placed into the same class as any TAL effector for which sequence data was available (maximum distance ≤ 5.0 and *p* ≤ 0.01). An alignment of the RVDs with other TAL effectors found in bacterial spot xanthomonads showed that the *avrBs3*-like effector shared several blocks of homology with *avrBs4*; however, differed by an average pairwise distance of 8.8 RVDs. Inoculation of strains AL37 and AL57 on tomato cv. Bonny Best and MoneyMaker, with the *Bs4* resistance gene, resulted in a compatible interaction and indicated that this TALE was not recognized by *Bs4* (Fig. 2).

**Figure 2.**
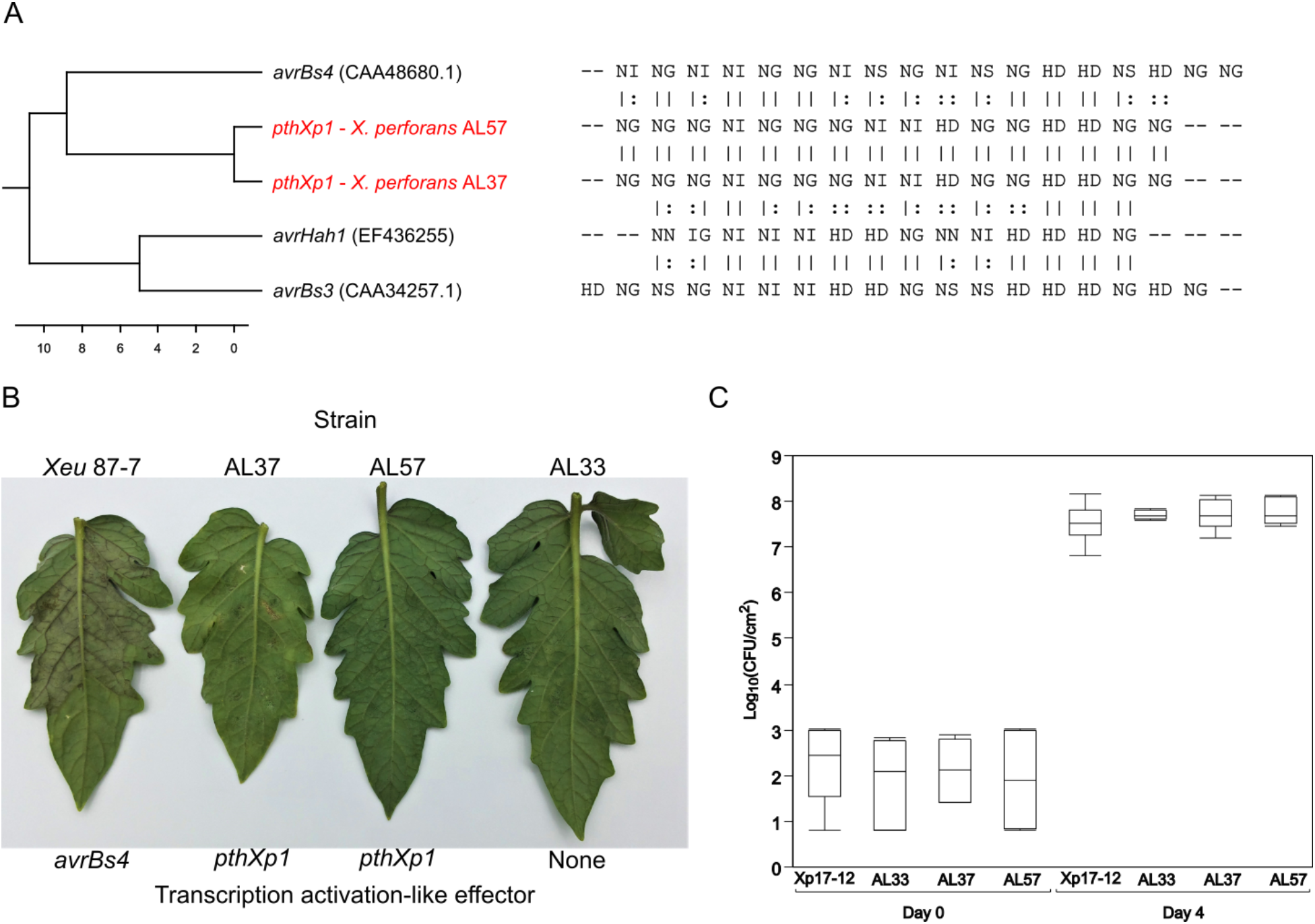
Identification of a new class of transcription activation-like effector, *pthXp1*, in *X. perforans*. (A) Dendrogram (left) constructed from an alignment of repeat variable di-residues (right) found among the *avrBs3* family of effectors previously described in bacterial spot causing xanthomonads and *pthXp1* from *X. perforans* strains AL37 and AL57 (shown in red). The scale bars indicate the number of substitutions per site. (B) Differential reactions of *X. perforans* strains with and without *pthXp1* and *X. euvesicatoria* strain 87-7 with *avrBs4* in tomato cv. MoneyMaker which has *Bs4* resistance. Tomato leaves were infiltrated with a bacterial suspension raised to 10^8^ CFU ml^−1^. A hypersensitive resistance is indicated by the appearance of collapsed/necrotic leaf tissue, 24h after inoculation. (C) Bacterial growth of *X. perforans* strains with (AL37 and AL57) and without (Xp17-12 and AL33) *pthXp1*, in tomato cv. BonnyBest. Plants were infiltrated with a bacterial suspension raised to 10^4^ CFU ml^−1^. Four replications were included for each treatment and the experiment was carried out twice with similar results. No significant differences in growth were observed four days after inoculation (*P* < 0.05).

### Novel diversity within *Xp* emerged through recent recombination derived from outside of the bacterial spot species complex

To examine patterns of homologous recombination between *Xp*, *Xeu*, and other closely related *Xanthomonas* spp., a second core genome alignment was constructed utilizing the strains sequenced in this study and respective genome assemblies available from GenBank (*n* = 68). A maximum likelihood phylogeny generated from the resulting 3.98 Mb alignment was largely congruent to the population structure inferred by previous studies (18, 20) and showed *Xp* and *Xeu* branching from each other into two distinct phylogenetic groups, while other related *Xanthomonas* spp. displayed considerably longer branch lengths and clustered into a paraphyletic group of strains. Analysis with FastGear software indicated the presence of three distinct lineages, which were defined as a group of strains that shared a common ancestry in at least half of the alignment (3) and corresponded to the three phylogenetic groups described above (Fig. 3A).

**Figure 3.**
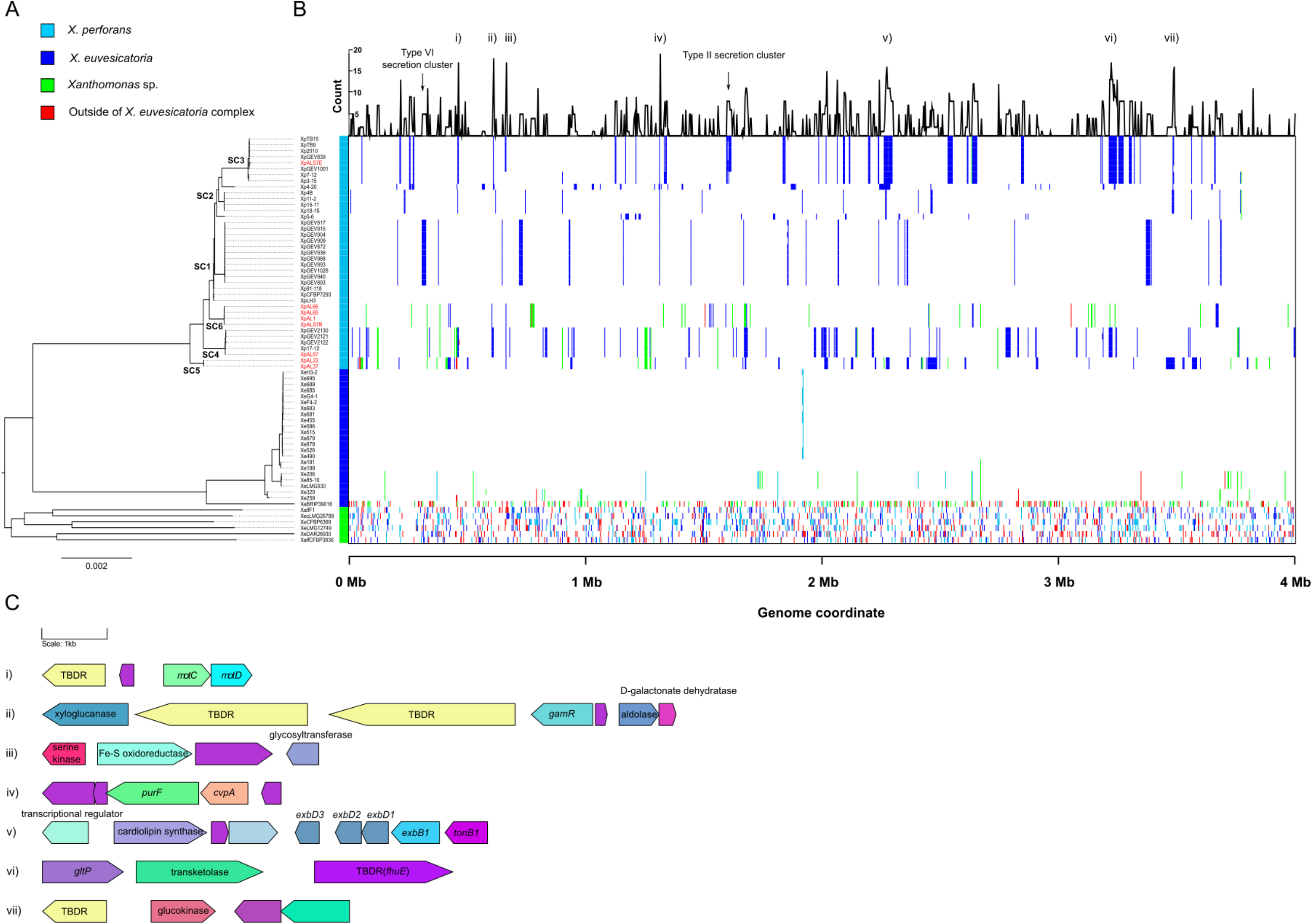
Landscape of homologous recombination across the *X. euvesicatoria* species complex. (A) Phylogeny of 67 *X. perforans*, *X. euvesicatoria*, and strains of related *Xanthomonas* species based on a core genome alignment of 3.98 Mb. The sequence clusters inferred within *X. perforans* (see Figure 1) are labeled at the nodes and the strains sequenced in this study are highlighted in red. Lineages predicted by FastGear software are color coded to the right of tree. (B) Patterns of recent recombination across the core genome of *X. perforans*, *X. euvesicatoria*, and related *Xanthomonas* species. Colors indicate donor lineage of the recent recombination events and the line graph above shows the frequency of recombination within *X. perforans* at a particular site in the alignment. Recombination events originating from outside of the sampled *Xanthomonas* population are shown in red. (C) Gene content of core genome loci with elevated recombination frequency (numbered i to vii) within *X. perforans*. Genes are color coded-coded according to their annotation, with hypothetical proteins shown in purple. Gene names/functional annotations are shown where available. The abbreviation, TBDR, indicates a putative TonB-dependent receptor.

A total of 3,174 recent recombination events were identified across the *Xeu* species complex (Fig. 3B). Consistent with the long branch lengths, the lineage composed of various *Xanthomonas* spp. displayed evidence of extensive recent recombination, with an average (± standard deviation) 19.00 ± 0.88% of the core genome predicted to be recombinant among the individual strains. A considerable amount of recent recombination found within this group originated from both the *Xp* and *Xeu* lineages (7.31 and 5.90% of the alignment respectively); however, the proportion of gene flux from each donor lineage was somewhat variable at the strain level (standard deviations of ± 2.16 and ± 1.19% for *Xp* and *Xeu* respectively). Interestingly, approximately a third of the recombinant sequences detected in this lineage (5.90 ± 2.33% of the alignment) were predicted to have originated from outside of the sampled *Xanthomonas* population (Fig. 4).

**Figure 4.**
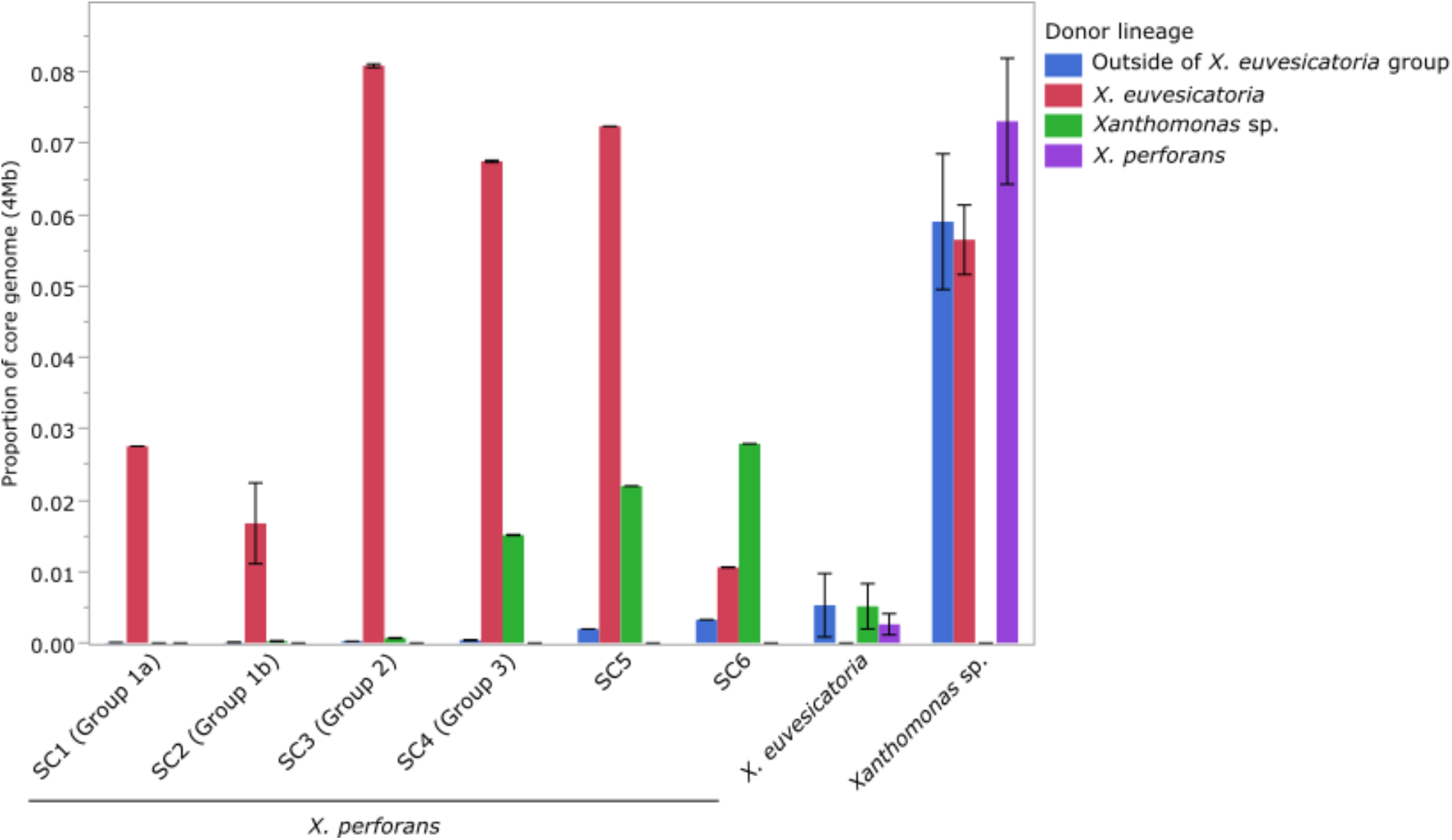
Summary of the proportion and origin of recent recombination among lineages within *X. euvesicatoria* species complex, as predicted by FastGear. The phylogroup designation of the *X. perforans* sequence clusters (SCs) as described in previous studies are shown in parentheses (19, 20). Outside of *X. euvesicatoria* group shows the proportion of recombinant sequences donated from outside of the sampled *Xanthomonas* population. The error bar indicates that standard deviation of the mean.

Evidence of recent recombination was also apparent within the *Xp* lineage. Nearly all of the recent recombination events detected in the Florida *Xp* strains collected prior to 2015 (SCs 1 through 3) were predicted to have originated from *Xeu*, whereas recombination events derived from both *Xeu* and the lineage composed of various *Xanthomonas* spp. were detected in the recently described, Group 3 *Xp* strains (designated here as SC4), as well as the Alabama strains located in SCs 5 and 6. The average proportion of core genome donated from various *Xanthomonas* spp. ranged from 1.51 to 2.79% among these groups and was the primary donor of recent recombination (66.7% of total recent recombinant sequences) to the pepper pathogenic strains of SC6. Variability in the overall proportion of recent recombination was also observed among the *Xp* sequence clusters ranging from 1.41 ± 1.70% in SC1 to 9.64 ± 0.00% in SC5 (Fig. 4).

### Recombination hot-spots are characterized by pathways involved in the acquisition/metabolism of plant-derived nutrients and motility

We investigated seven recombination “hot-spots” within *X. perforans* based on the frequency of recombination events at a particular site in the core genome alignment (Fig. 3C). Hot-spots (ii), (vi), and (vii) mapped to loci with genes involved in carbohydrate and amino acid metabolism including a xyloglucanase, *lysR*-type regulator of galactose metabolism (*gamR*), transketolase, a symporter of protons/glutamate (*gltP*), and a glucokinase. All of these genes were adjacent to at least one TonB-dependent receptor (TBDR) while a fourth TBDR locus, hot-spot (i) contained the flagellar motor components *motA* and *motB.* It was therefore interesting to find the entire TonB-transduction system (*tonB*-*exbB*-*exbD1*-*exbD2*) at recombination hot-spot (v). Other genes in recombination hot-spots included an iron-sulfur oxidoreductase (iii) and a gene with homology to colicin V production protein *cvpA* (iv). Finally, while not necessarily present in hot-spots, we noted that most of the genes which encode for the Xps type II and type VI secretion systems were recombinant among strains from SCs 3 and 1, respectively.

## Discussion

Upon introduction of *X. perforans* (*Xp*) to Florida tomato fields in 1991, the bacterial species has undergone several changes over the past two decades including overtaking *X. euvesicatoria* (*Xeu*) as the predominant pathogen of tomato, a shift in race associated with null mutations affecting the type 3 secreted effector *avrXv3*, and divergence into three different phylogenetic groups (12, 19, 20, 25). Until the isolation of a single *Xp* strain from an infected pepper sample during the 2010 season in Florida (20), the host-range of this bacterial pathogen was thought to be restricted to tomato. Here, we report for the first time the genomes of several *Xp* strains that were isolated from naturally infected pepper plants grown in Alabama as well as four strains isolated from tomatoes in the state (Table 1).

**Table 1.** Collection information, assembly statistics, and pathogenicity phenotyping on differential pepper varieties for the *X. perforans* strains sequenced in this study.

| Strain | Host | County | Contigs (N) | N50 (kb) | Genome size (Mb) | GenBank Accession No. |
| --- | --- | --- | --- | --- | --- | --- |
| AL1 | Tomato | Lee | 86 | 1.78 | 5.02 | SMVJ00000000 |
| AL65 | Pepper | Lee | 58 | 1.91 | 5.02 | SMVI00000000 |
| AL66 | Pepper | Lee | 52 | 2.49 | 5.02 | SMVH00000000 |
| AL57 | Tomato | Lee | 55 | 2.68 | 4.97 | SMVC00000000 |
| AL33 | Tomato | Tuscaloosa | 60 | 2.27 | 5.22 | SMVE00000000 |
| AL37 | Tomato | Tuscaloosa | 98 | 1.31 | 5.28 | SMVD00000000 |
| ALS7B | Pepper | Tuscaloosa | 65 | 1.87 | 5.06 | SMVG00000000 |
| ALS7E | Pepper | Tuscaloosa | 88 | 1.39 | 5.15 | SMVF00000000 |

Maximum likelihood reconstruction of the *Xp* core genome revealed the presence of novel diversity among the strains sequenced in this study, with the identification of two additional phylogenetic groups, designated here as sequence clusters (SCs) 5 and 6 (Fig. 1). To our surprise, the majority of pepper strains did not group with others within SC3 that were previously found to be pathogenic to pepper but comprised the newly emerged SC6. Three of the four strains located within this phylogenetic group carried a plasmid identical to one often found in *X. gardneri*, which harbored two different type 3 secreted effectors including *xopAO* and the transcription activation-like effector (TALE) *avrHah1* (Table 2). The latter effector is commonly associated with enhanced virulence to pepper plants through hijacking the host-expression of pectate lyase genes, which results in a profuse water-soaking phenotype and proliferation of bacterial growth (26, 27). However, as the pepper strain ALS7B lacked the plasmid associated with the horizontal transfer of *avrHah1*, it is clear that this TALE contributes to increased symptom development but is not necessary for pathogenicity to pepper among strains of this genetic background (Fig. S1). Interestingly, strain ALS7B was isolated from plants collected from same pepper field in Tuscaloosa county as ALS7E of SC3, indicating the occurrence of mixed infections among *Xp* lineages.

A second unique phylogenetic group, referred to here as SC5, was identified in our analyses and comprised two Alabama strains that were isolated from tomato plants collected from commercial production fields. This phylogenetic group branched separately from SC4 (also referred to as group 3), which was recently found to be prevalent in Florida tomato fields (19). Screening of the genome assemblies with tBLASTn revealed that the Alabama strains located in these two sequence clusters carried an intact copy of the avirulence gene *avrXv3*, which as has not been observed in the *Xp* population surveyed in the Florida since 2006 (14, 20). Interestingly, the *avrXv3* allele found in the strains sequenced here differed from other available sequences by two amino substitutions located at both the N- and C-terminal domains of the protein (data not shown); however, these mutations did not allow for *avrXv3* to escape host-recognition (Table 2).

**Table 2.**
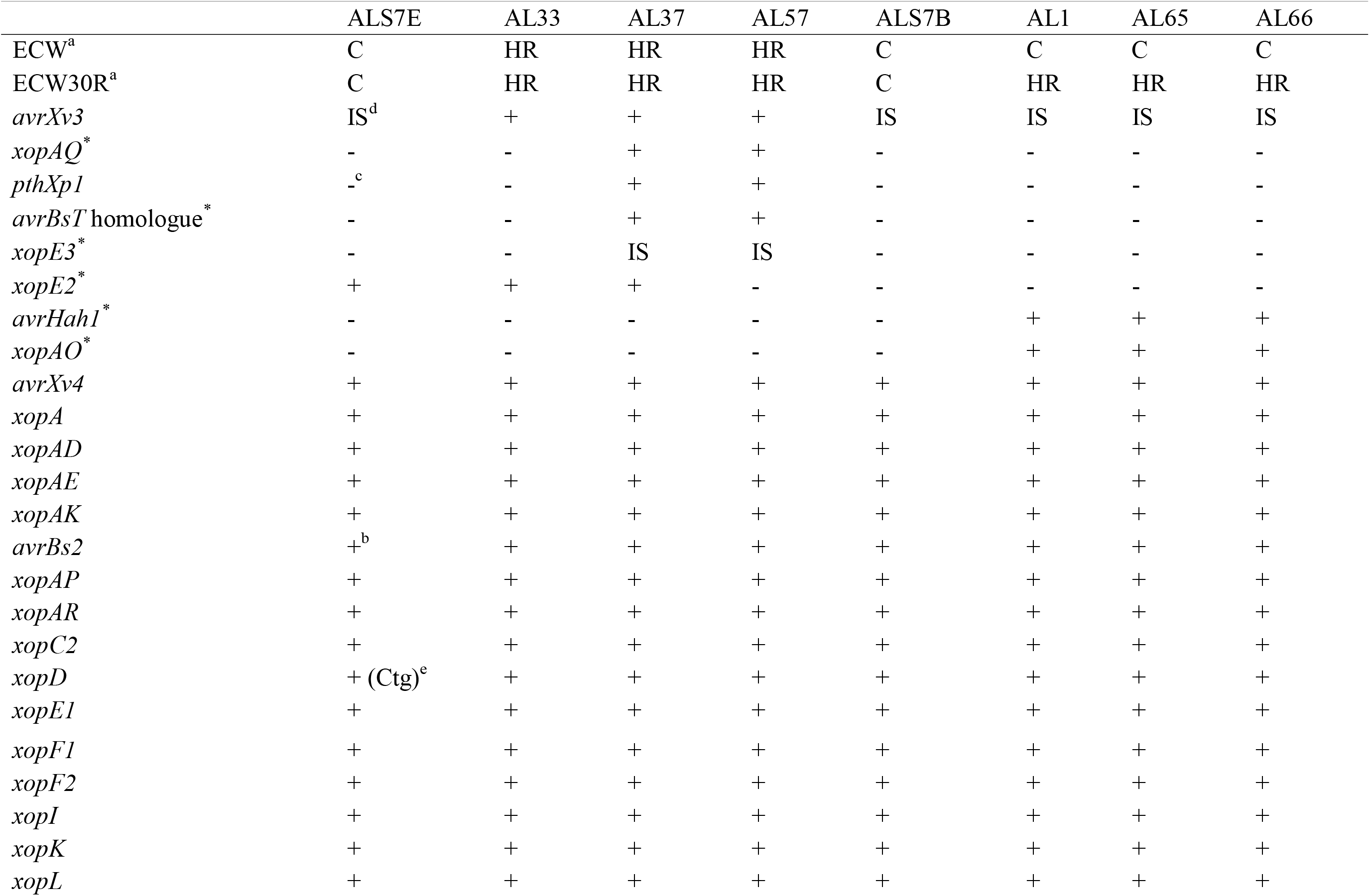
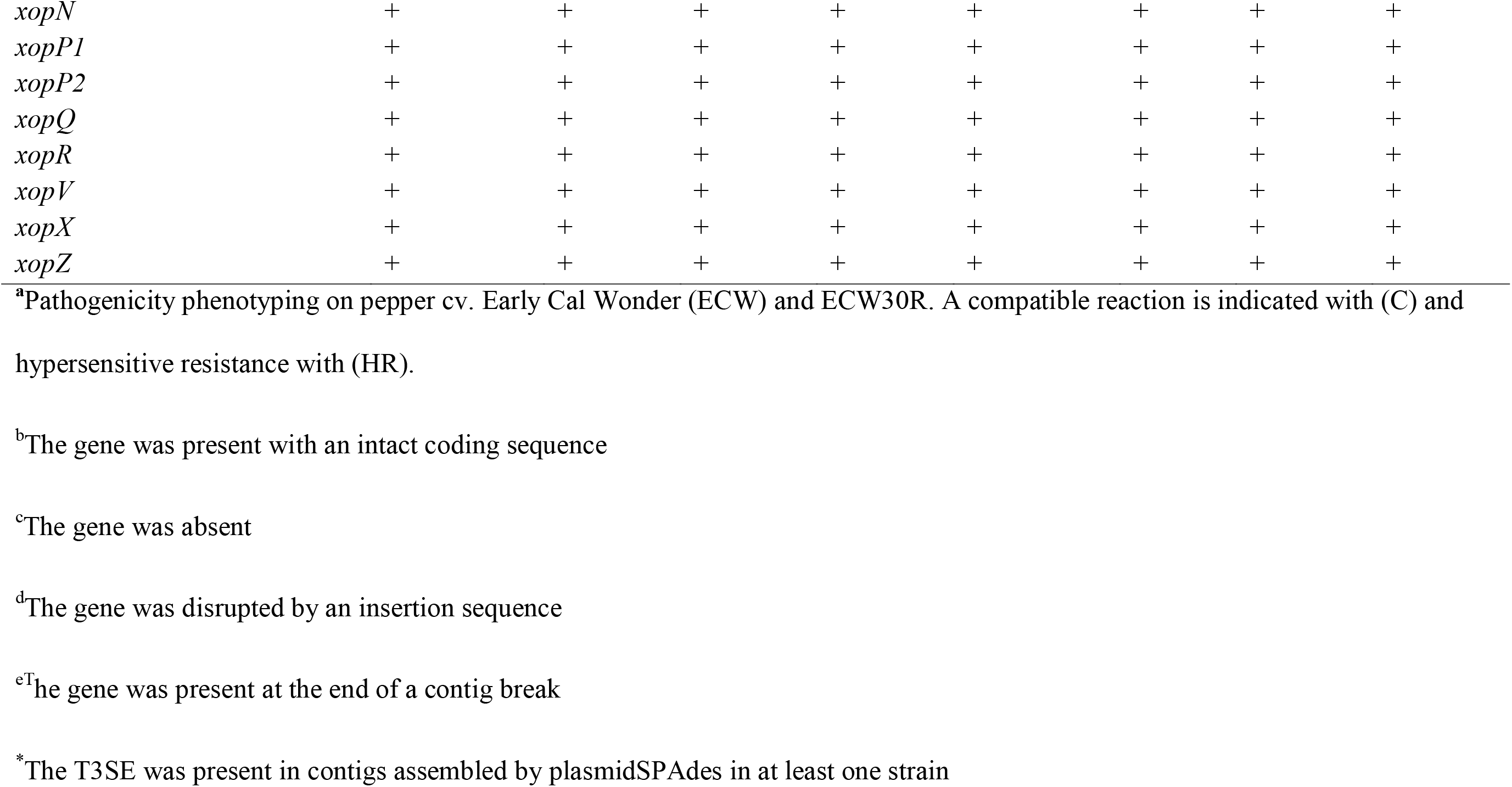
Pathogenicity phenotyping and distribution of type 3 secreted effectors among *X. perforans* strains isolated from tomato and pepper in Alabama.

Strains AL57 and AL37, of SCs 4 and 5 respectively, also carried a TALE which belonged to the *avrBs3*/*avrBs4* family of effectors commonly found in *X. euvesicatoria* and *X. vesicatoria*. To our knowledge, this is the first identification of an *avrBs3-*like in *Xp.* Moreover, it was surprising to find an effector from this family in strains isolated from tomato as *avrBs4* serves an avirulence gene when expressed in plants with the cognate *Bs4* resistance gene, while *avrBs3* does not elicit a hypersensitive response, it is likely a negative factor (28). We confirmed that the *Xp* strains carrying this effector did not induce a *Bs4*-mediated resistance in tomato cv. Moneymaker and grew to high populations, comparable to strains devoid of TALEs, in tomato cv. Bonny Best (Fig 2). Analysis of the repeat variable di-residues supported these observations and showed that the predicted binding specificity of the gene was divergent from other TALEs recorded in *Xanthomonas* spp., suggesting the evolution of this effector to avoid *Bs4*-mediated recognition. As the gene could not be assigned to the same class as any TALE for which sequence data was available, we propose the name *pthXp1.* Interestingly, we also noted evidence of a similar *avrBs3*-like effector in the genome assemblies of the Florida *Xp* strains located in SC4 and therefore, may be a significant factor contributing to the recent emergence of this pathogen lineage (data not shown).

Investigation of these strains in the context of the larger *Xeu* species complex revealed a dynamic pattern of gene flow between *Xp*, *Xeu*, and a third lineage which was composed of various *Xanthomonas* spp. (Fig. 3). The latter lineage exhibited evidence of extensive admixture, with ~19% of the core-genome affected by recent recombination events derived from both *Xp, Xeu*, and an unidentified donor outside of the sampled *Xanthomonas* population (Fig. 4). As these strains were isolated from numerous hosts including pepper, alfalfa, citrus, onion, Anthurium, and Commiphora, among others (Table S1), this could be an indication of a cryptic ecology within the *Xeu* species complex. Host-range studies of *Xeu* and related pathogens have shown that strains often have the capacity to colonize and/or infect several plant species beyond their original host of isolation (29–32), thus supporting this hypothesis. However, because the individual branches found within this apparently host-generalist lineage were as equally diverged as that of the *Xp* and *Xeu* lineages (Fig. 3A), it is possible that these contrasting evolutionary patterns may reflect adaptation to diverse hosts.

Consistent with previous observations (19), *Xeu* was the primary donor of recent recombination to *Xp* (Fig 3B). Each of the SCs inferred within this lineage were affected by recent recombination to varying degrees (Fig. 4), which correlated with the branch lengths observed in the core-genome phylogeny (Fig 1). Interestingly, the recent emergence of SCs 4, 5, and 6 was associated in-part with the acquisition of recombinant sequences from the lineage composed of various *Xanthomonas* spp., which were absent from the *Xp* strains found to predominant in Florida prior to 2015 (SCs 1 through 3; Fig. 4). Surprisingly, this lineage was the primary donor of recent recombination to the pepper pathogenic strains of SC6, for which only a minor proportion of the core genome (4.18%) carried signals of homologous recombination. This contrasted with the pepper pathogenic strains of SC3, which acquired ~8.10% of the core-genome from *Xeu* (Fig. 4) and suggests that the evolution of pathogenicity to pepper among strains from these two phylogenetic groups has likely emerged independently.

Examination of recombination hot-spots within *Xp* revealed that several loci implicated in the plant-pathogen interaction were frequently recombining (Fig. 3C). Most of these recombination tracts contained genes associated with nutrient acquisition/metabolism (*gamR*, *gltP*, *fhuE*, a transketolase, and glucokinase) and motility (*motC*, *motD*), and were often adjacent to at least one TonB-dependent receptor. This class of outer membrane receptor binds with high specificity to a variety of macromolecules, including plant-derived carbohydrates and iron, and facilitates the active transport of substrates into the periplasm via the TonB transduction system (33). It was therefore curious to note that the core components of this system (*tonB1, exbB1*, *exbD1*, *exbD2*, and *exbD3*) were also present in a separate recombination hot-spot. One gene, *exbD2*, distinguished the TonB operon found in *Xanthomonas* spp. from that of most other bacterial genera and is essential for pathogenicity and induction of bacterial pectate lyase activity (34, 35). Interestingly, we found that in addition to the TonB system, the entire Xps type II secretion cluster was recombinant among the pepper pathogenic strains of SC3. The importance of the Xps type II gene cluster in the secretion of numerous plant-cell wall degrading enzymes and virulence has been well established in *Xeu* (36–38). Therefore, it is possible that the recombination-mediated remodeling of the TonB and Xps type II systems may affect the secretion of plant cell-wall degrading enzymes and the regulation of downstream pathogenesis processes, leading to adaptation within the pepper niche.

Further work is required to test this hypothesis and to investigate the evolutionary processes enabling certain *Xp* lineages to infect pepper plants in the absence of a gene for gene interaction, while the host-range of others remains limited to tomato (20). Given the population structure identified here, genome-wide association analysis may be an appropriate tool to investigate this phenomenon more rigorously. Overall, the diversity of the *Xp* population observed in Alabama was striking in relation to that reported in neighboring tomato and pepper production regions in the United States and was indicative of an adaptive evolution. Recently, TALEs were also identified in *Xp* strains collected in Louisiana, USA and in Italy (Jones, *unpublished data*), pointing towards the recent acquisition of these virulence factors as a trend extending beyond the *Xp* population sampled in Alabama. Taken together, the results of this study highlight the need for regular pathogen surveillance when selecting gene candidates for resistance breeding.

## Material and Methods

### Bacterial strain collection, sequencing, and assembly

Eight bacterial strains, isolated from symptomatic tomato and pepper plants, grown in three different Alabama counties during the summer of 2017 (Table 1), were selected at random for draft genome sequencing. Genomic DNA was isolated using the CTAB-NaCl method, as described previously (39), and submitted to the Georgia Genomics and Bioinformatics Core, University of Georgia for library preparation and sequencing. Paired-end reads were generated by multiplexing 12 libraries in a single lane on an Illumina MiSeq Micro (PE150) and *de novo* assemblies were constructed using the A5-miseq pipeline under the default settings (v.20160825, 40). Briefly, adapter and quality trimming were performed with Trimmomatic (41), followed by error correction with the kmer-based String Graph Assembler algorithm (42), and contig assembly using the Iterative de Bruijn Graph Assembler (43). The genome assemblies and raw sequencing reads were submitted to NCBI GenBank under BioProject accession number PRJNA526717.

### Reconstruction of the *X. perforans* core genome

To test for a nested population structure within *X. perforans*, a read-mapping approach was taken using the Snippy pipeline (v.4.3.5, 44). The sequencing reads for 33 *X. perforans* strains described by Schwartz et al. (20) and Timilsina et al. (19) were downloaded from the Sequence Read Archive database (PRJNA526741) and subjected to adapter and quality trimming with Scythe (v.0.991, https://github.com/vsbuffalo/scythe) and SolexaQA respectively (v.3.1.7.1, 45). These data were combined with the quality-trimmed reads generated in this study and were individually aligned against the completed genome of *X. perforans* strain 91-118 using the Burrows-Wheeler Alignment tool (v.0.7.17-r1188, 46). Only reads with a mapping quality of 60 (i.e., uniquely mapped reads) and Phred quality score of 20 (at least 99% consensus) were included in the analysis, while soft-clipped reads were filtered from the data-sets with SAMtools (v.1.9, 47). Variants were called from the whole genome alignments using FreeBayes (v.1.2.0, 48) and single-nucleotide polymorphisms (SNPs) common to all genomes were extracted to generate a concatenated set of high-quality core-genome SNPs. This concatenated SNP alignment was used to infer the population structure of *X. perforans* using HierBAPS software with four hierarchy levels and an upper cluster limit of 20 (24). Additionally, a maximum likelihood phylogeny was inferred from the concatenated SNP alignment using iQTree (v.1.6.4, 49). The best fitting model (TVM+ASC+G4) was chosen using Model Finder (50) and branch support was assessed with the ultrafast bootstrap method using 1000 replicates (51). The phylogenetic tree was visualized and annotated using FigTree (v.1.4.2, http://tree.bio.ed.ac.uk/software/figtree/).

### Interlineage recombination

To examine patterns of gene flow across the *X. euvesicatoria* species complex as described by Parkinson et al. (23), a core genome alignment was generated with the program Parsnp (52) using the genome assemblies of 39 *X. perforans* strains, 23 of *X. euvesicatoria*, and six of related *Xanthomonas* spp. available from GenBank (Table S1). Highly fragmented genome assemblies (≥ 400 contigs) were excluded from this analysis as they were found to significantly reduce the alignment coverage of the reference genome. The xmfa alignment file produced by Parsnp was converted to a multi-fasta format with the perl script xmfa2fasta.pl (https://github.com/kjolley/seq_scripts/blob/master/xmfa2fasta.pl) and used as input for the FastGear algorithm (3). This software was run under default settings with the statistical significance of recombination predictions tested using a Bayes factor (BF) > 1 for recent recombination events and BF > 10 for ancestral recombination events. The resulting output was visualized using the Phandango web-server (53) and recombination “hot-spots” within *X. perforans* were visually assessed based on the frequency of recombination events at a particular site in the alignment after correcting for oversampling of clonal populations. Gene neighborhoods located in recombination hot-spots were extracted for the 91-118 reference chromosome and visualized using Gene Graphics (54). Finally, the core genome alignment was used to construct a maximum likelihood phylogeny using RAxML (v8.2.10, 55) with the Gamma Time Reversible model of nucleotide substitution and 1000 bootstrap replicates.

### Prediction of type 3 secreted effectors (T3SEs) and plasmids

A database of T3SEs was compiled using the *Xanthomonas* web resource (http://www.xanthomonas.org/t3e.html) and used to query the genome assemblies generated in this study using tBLASTn (E-value ≤ 10^−5^). An effector was considered to be present if it displayed ≥ 80% aa over 80% of the query length. Nucleotide sequences of putative effector genes were subsequently extracted from the genomes and examined for frameshift mutations and other potential disruption of the coding sequence with BLASTx. Plasmids were computationally predicted using plasmidSPAdes software (56). Contigs assembled by the program were subsequently screened against the NCBI nucleotide collection database (nr/nt) to assess for the presence of putative plasmid sequences. These contigs were also screened for T3SEs as described above.

### Sequencing and analysis of *pthXp1* from *X. perforans* strains AL37 and AL57

After noting evidence an *avrBs3*-like effector in the genome assemblies of strains AL37 and AL57, forward (5’–ATGAGGTGCAATCGGGTCTG-3’) and reverse (5’– GTCCTCATCTTGTTCCCGCA-3’) primers were designed to anneal to conserved loci within the N- and C-terminal domains of *avrBs3*. Phusion high fidelity polymerase chain reaction (PCR) was used to amplify the gene with a T100 Thermal Cycler (BioRad, Hercules, CA). Each PCR reaction contained in a final volume of 50 µl with 1 x Phusion Master Mix containing HF buffer (Thermo Scientific, Waltham, MA), 0.5 µM of each primer, and 1 µl of cells treated at 95□ for 10 min. The cycling conditions consisted of an initial denaturation at 98°C for 30 s, followed by 30 cycles of denaturation at 98°C for 10 s, annealing at 63°C for 30 s, extension at 72°C for 90 s, and a final extension of 72°C for 10 min. The resulting amplicons were purified and submitted to Eurofins Genomics (Louisville, KY) for sanger sequence using the Power Read Service optimized for tandem repeat stretches with the internal sequencing primers (5’– AAGATTGCAAAACGTGGCGG-3’) and (5’–CCGGATCAGGGCGAGATAAC-3’). Classification of the completed transcription activation-like effector sequences and alignment of the repeat variable di-residues were conducted using AnnoTALE (v.1.4) software utilizing the default alignment parameters (57). The completed *pthXp1* sequences were submitted the NCBI GenBank with the accession numbers MK755838 and MK755839.

### Plant inoculations

The strains sequenced in this study were assessed for the capacity to induce a hypersensitive resistance in three to four-week old pepper cv. Early California Wonder (ECW) and ECW30R as well as tomato cv. BonnyBest and MoneyMaker. Individual leaves were infiltrated with a needless syringe using a bacterial suspension raised to 10^8^ CFU ml^−1^ (OD_600nm_ = 0.3) in a sterile MgSO_4_ * 7H_2_O solution. The presence of necrotic, collapsed tissue, 24h after inoculation was scored as a positive result. The pathogenicity of strains was assessed using bacterial suspension prepared as described above and diluted to a concentration of 10^4^ CFU ml^−1^ using 10-fold serial dilutions. Bacterial populations were enumerated at 0 and 4 days after inoculation as described by Schwartz et al. (20) and a two-way analysis of variance was conducted to test for differences between treatments using JMP Pro 13 software (SAS Institute, Cary, NC). All plants were kept under standard greenhouse conditions and each strain was tested for the reactions described above at least twice.

## Acknowledgements

This work was supported by the USDA National Institute of Food and Agriculture, Hatch project 1012760 and Alabama Agricultural Experiment Station. We thank Alabama Cooperative Extension agents for their support in the collection of samples from the fields.

