## Supplemental material for "Recently emerged and diverse lineages of *Xanthomonas perforans* have independently evolved through plasmid acquisition and homologous recombination originating from multiple *Xanthomonas* species"

| **Table S1**. Collection information and GenBank accession number for *Xanthomonas* species used in comparative analysis. | | | | |
| --- | --- | --- | --- | --- |
| Organism/Name | Strain | BioProject | Assembly | Host |
| *X. perforans* | LH3 | PRJNA344031 | GCA_001908855.1 | Tomato |
| *X. perforans* | Xp5-6 | PRJNA276609 | GCA_001009365.1 | Tomato |
| *X. perforans* | Xp7-12 | PRJNA276610 | GCA_001009385.1 | Tomato |
| *X. perforans* | Xp10-13 | PRJNA276613 | GCA_001009405.1 | Tomato |
| *X. perforans* | Xp11-2 | PRJNA276614 | GCA_001009445.1 | Tomato |
| *X. perforans* | Xp15-11 | PRJNA276615 | GCA_001009465.1 | Tomato |
| *X. perforans* | GEV839 | PRJNA276632 | GCA_001009475.1 | Tomato |
| *X. perforans* | Xp3-15 | PRJNA276600 | GCA_001009675.1 | Tomato |
| *X. perforans* | Xp4-20 | PRJNA276608 | GCA_001009705.1 | Tomato |
| *X. perforans* | Xp17-12 | PRJNA276616 | GCA_001009745.1 | Tomato |
| *X. perforans* | Xp18-15 | PRJNA276617 | GCA_001009765.1 | Tomato |
| *X. perforans* | GEV909 | PRJNA276640 | GCA_001009825.1 | Tomato |
| *X. perforans* | GEV940 | PRJNA276648 | GCA_001009885.1 | Tomato |
| *X. perforans* | GEV1001 | PRJNA276679 | GCA_001010025.1 | Tomato |
| *X. perforans* | CFBP 7293 | PRJNA350372 | GCA_001976075.1 | Tomato |
| *X. perforans* | GEV872 | PRJNA276635 | GCA_001009485.1 | Tomato |
| *X. perforans* | GEV893 | PRJNA276636 | GCA_001009545.1 | Tomato |
| *X. perforans* | Xp4B | PRJNA276599 | GCA_001009665.1 | Tomato |
| *X. perforans* | GEV904 | PRJNA276639 | GCA_001009795.1 | Tomato |
| *X. perforans* | GEV936 | PRJNA276644 | GCA_001009845.1 | Tomato |
| *X. perforans* | GEV915 | PRJNA276641 | GCA_001009855.1 | Tomato |
| *X. perforans* | GEV917 | PRJNA276642 | GCA_001009865.1 | Tomato |
| *X. perforans* | TB6 | PRJNA276688 | GCA_001009945.1 | Tomato |
| *X. perforans* | TB9 | PRJNA276689 | GCA_001009955.1 | Tomato |
| *X. perforans* | GEV993 | PRJNA276677 | GCA_001010005.1 | Tomato |
| *X. perforans* | GEV1026 | PRJNA276680 | GCA_001010035.1 | Tomato |
| *X. perforans* | TB15 | PRJNA276690 | GCA_001010105.1 | Tomato |
| *X. perforans* | Xp2010 | PRJNA276618 | GCA_001009785.1 | Pepper |
| *X. perforans* | GEV2121 | PRJNA436012 | GCA_004102075.1 | Tomato |
| *X. perforans* | GEV2122 | PRJNA436012 | GCA_004102435.1 | Tomato |
| *X. perforans* | GEV2130 | PRJNA436012 | GCA_004102405.1 | Tomato |
| *X. euvesicatoria* | 85-10 | PRJNA341901 | GCA_001854165.1 | Pepper |
| *X. euvesicatoria* | LMG930 | PRJNA344031 | GCA_001908795.1 | Pepper |
| *X. euvesicatoria* | 329 | PRJNA275489 | GCA_001008805.1 | Pepper |
| *X. euvesicatoria* | 206 | PRJNA275486 | GCA_001008815.1 | Pepper |
| *X. euvesicatoria* | 515 | PRJNA275494 | GCA_001008825.1 | Pepper |
| *X. euvesicatoria* | 526 | PRJNA275495 | GCA_001008835.1 | Pepper |
| *X. euvesicatoria* | 586 | PRJNA275496 | GCA_001008885.1 | Pepper |
| *X. euvesicatoria* | 679 | PRJNA275498 | GCA_001008895.1 | Pepper |
| *X. euvesicatoria* | 681 | PRJNA275499 | GCA_001008905.1 | Pepper |
| *X. euvesicatoria* | 678 | PRJNA275497 | GCA_001008915.1 | Pepper |
| *X. euvesicatoria* | 259 | PRJNA275487 | GCA_001008965.1 | Pepper |
| *X. euvesicatoria* | 199 | PRJNA275485 | GCA_001008975.1 | Pepper |
| *X. euvesicatoria* | 354 | PRJNA275490 | GCA_001008995.1 | Pepper |
| *X. euvesicatoria* | 683 | PRJNA275500 | GCA_001009095.1 | Pepper |
| *X. euvesicatoria* | 684 | PRJNA275501 | GCA_001009125.1 | Pepper |
| *X. euvesicatoria* | 685 | PRJNA275503 | GCA_001009135.1 | Pepper |
| *X. euvesicatoria* | F4-2 | PRJNA275506 | GCA_001009165.1 | Pepper |
| *X. euvesicatoria* | H3-2 | PRJNA275508 | GCA_001009175.1 | Pepper |
| *X. euvesicatoria* | 689 | PRJNA275504 | GCA_001009205.1 | Pepper |
| *X. euvesicatoria* | 695 | PRJNA275505 | GCA_001009215.1 | Pepper |
| *X. euvesicatoria* | G4-1 | PRJNA275507 | GCA_001009245.1 | Pepper |
| *X. euvesicatoria* | 181 | PRJNA275484 | GCA_001010095.1 | Pepper |
| *X. euvesicatoria* | BRIP39016 | PRJNA454505 | GCA_003993195.1 | Pepper |
| *X. axonopodis* pv. *citrumelo* | F1 | PRJNA62495 | GCA_000225915.1 | Citrus |
| *X. euvesicatoria* pv. *alfalfae* | CFBP 3836 | PRJNA212245 | GCA_000488955.1 | Alfalfa |
| *X. euvesicatoria* pv. *allii* | CFBP 6369 | PRJNA231977 | GCA_000730305.1 | Onion |
| *X. euvesicatoria* | LMG 12749 | PRJNA254447 | GCA_001401675.2 | Anthurium |
| *X. euvesicatoria* pv. *commiphorae* | LMG 26789 | PRJNA298623 | GCA_003698225.1 | Commiphora wightii |
| *X. euvesicatoria* | DAR26930 | PRJNA454505 | GCA_003993445.1 | Pepper |

| **Table S2**. Assembly statistics and BLAST results for the *X. perforans* strains sequenced in this study using plasmidSpades. | | | | | | |
| --- | --- | --- | --- | --- | --- | --- |
| Strain | Size | Coverage | Best Hit | Query Coverage | Identity | Notes |
| AL1 | 51220 | 269.654 | Uncultured bacterium plasmid pAKD18 | 100% | 99% | Broad host-range plasmid (soil) |
|  | 18716 | 350.07 | Xanthomonas gardneri strain JS749-3 plasmid pJS749-3.2 | 100% | 100% |  |
|  | 13436 | 314.922 | Xanthomonas gardneri strain JS749-3 plasmid pJS749-3.2 | 100% | 100% |  |
|  | 7032 | 286.607 | Xanthomonas gardneri strain JS749-3 plasmid pJS749-3.2 | 100% | 100% |  |
|  | 2862 | 349.353 | Xanthomonas gardneri strain JS749-3 plasmid pJS749-3.2 | 100% | 99% |  |
|  | 1513 | 321.85 | Xanthomonas gardneri strain JS749-3 plasmid pJS749-3.2 | 100% | 100% | *avrHah1* |
| AL65 | 51220 | 41.9348 | Uncultured bacterium plasmid pAKD18 | 99% | 99% | Broad host-range plasmid (soil) |
|  | 36638 | 61.128 | X. gardneri JS749-3 plasmid pJS749-3.2 | 100% | 99% | *xopAO* |
|  | 8608 | 58.8971 | X. gardneri JS749-3 plasmid pJS749-3.2 | 100% | 99% | *avrHah1* |
| AL66 | 51220 | 51.9869 | Uncultured bacterium plasmid pAKD18 | 99% | 99% | Broad host-range plasmid (soil) |
|  | 21501 | 74.1444 | Xanthomonas gardneri strain JS749-3 plasmid pJS749-3.2 | 100% | 99% | *xopAO* |
|  | 20191 | 75.1247 | Xanthomonas gardneri strain JS749-3 plasmid pJS749-3.2 | 100% | 100% |  |
|  | 1601 | 78.8274 | Xanthomonas gardneri strain JS749-3 plasmid pJS749-3.2 | 100% | 100% | *avrHah1* |
| ALS7B | 34722 | 100.963 | Xanthomonas citri pv. citri strain TX160042 plasmid unnamed2 | 86% | 98% |  |
| AL37 | 26260 | 68.0367 | Xanthomonas hortorum strain B07-007 plasmid pB07007 | 88% | 91% |  |
|  | 17723 | 22.4369 | Xanthomonas campestris pv. campestris str. CN18 plasmid unnamed3 | 96% | 99% | *xopAQ* |
|  | 14092 | 23.3366 | Xanthomonas campestris pv. campestris str. CN03 plasmid unnamed | 90% | 99% |  |
|  | 6835 | 25.1465 | Xanthomonas arboricola pv. pruni str. CFBP 5530 plasmid pXap41 | 97% | 98% | *xopE3* |
|  | 5524 | 65.2752 | Xanthomonas citri pv. phaseoli var. fuscans strain CFBP6167 plasmid pC | 68% | 94% |  |
|  | 4650 | 21.1658 | Xanthomonas campestris pv. arecae strain NCPPB 2649 plasmid pXCARECAE29 | 87% | 94% | *avrBst* homologue |
| AL33 | 234562 | 15.129 | Xanthomonas gardneri strain ICMP7383 plasmid pICMP7383.1 | 77% | 98% | copper resistance,  pectate lyase,  *xopE2* |
|  | 20988 | 149.211 | Xanthomonas hortorum strain B07-007 plasmid pB07007 | 92% | 91% |  |
|  | 5524 | 142.155 | Xanthomonas citri pv. phaseoli var. fuscans strain CFBP6167 plasmid pC | 68% | 94% |  |
| AL57 | 34398 | 16.6196 | Xanthomonas campestris pv. campestris str. CN18 plasmid unnamed3 | 94% | 99% | *xopAQ, xopE3* |
|  | 6833 | 17.9698 | Xanthomonas arboricola pv. pruni str. CFBP 5530 plasmid pXap41 | 97% | 98% | *xopE3* |
| ALS7E | 42757 | 17.0581 | Xanthomonas perforans strain LH3 plasmid pLH3.2 | 97% | 99% |  |
|  | 12795 | 26.466 | Xanthomonas perforans strain LH3 plasmid pLH3.2 | 97% | 99% | copper resistance,  pectate lyase,  *xopE2* |

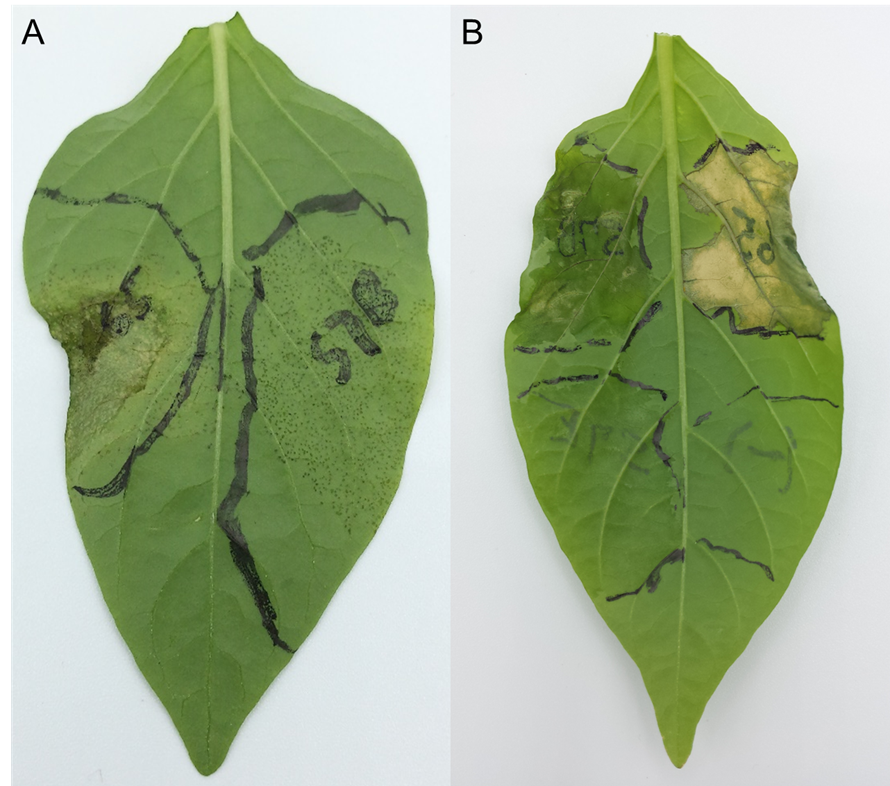

**Figure S1.** Pathogenicity phenotyping of *X. perforans* strains isolated from pepper. (A) Bacterial spot symptoms produced four days after inoculation by two *X. perforans* strains of sequence cluster (SC) 6, with (left) and without (right) the transcription-like effector *avrHah1*. Plants were infiltrated with a bacterial suspension raised to 10^4^ CFU ml^-1^. (B) Compatible (left) and incompatible (top right) interactions on pepper cv. ECW30R, shown 48 h after inoculation with 10^8^ CFU ml^-1^. A hypersensitive resistance mediated by the recognition of *avrHah1* in strain AL65 is shown on the top right portion of the leaf. Pepper strains ALS7B and ALS7E, from SCs 6 and 3 are shown on the top-left and bottom-left portion of the leaf respectively.

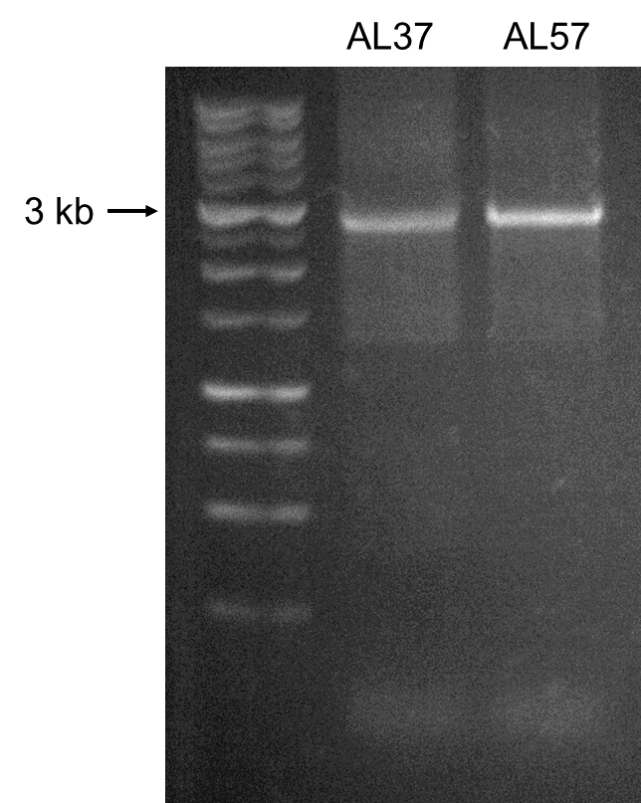
**Figure S2**. Amplicons of produced using primers designed to anneal to conserved loci within the N- and C-terminal domains of *avrBs3* family of transcription activation-like effectors for *X. perforans* strains AL37 and AL57.
